# DeepRibo: precise gene annotation of prokaryotes using deep learning and ribosome profiling data

**DOI:** 10.1101/317180

**Authors:** Jim Clauwaerts, Gerben Menschaert, Willem Waegeman

**Affiliations:** KERMIT, Department of Data Analysis and Mathematical Modelling, Ghent University, Coupure Links 653, 9000 Gent, Belgium; BioBix, Department of Data Analysis and Mathematical Modelling, Ghent University, Coupure Links 653, 9000 Gent, Belgium

## Abstract

Annotation of gene expression in prokaryotes often finds itself corrected due to small variations of the annotated gene regions observed between different (sub-species. It has become apparent that traditional sequence alignment algorithms, used for the curation of genomes, are not able to map the full complexity of the genomic landscape. We present DeepRibo, a novel neural network applying ribosome profiling data that shows to be a precise tool for the delineation and annotation of expressed genes in prokaryotes. The neural network combines recurrent memory cells and convolutional layers, adapting the information gained from both the high-throughput ribosome profiling data and Shine-Dalgarno region into one model. DeepRibo is designed as a single model trained on a variety of ribosome profiling experiments, and is therefore evaluated on independent datasets. Through extensive validation of the model, including the use of multiple species sequence similarity and mass spectrometry, the effectiveness of the model is highlighted.

## 1. Introduction

After more than 20 years of genome sequencing, it has become clear that the genomic diversity in bacteria is much larger than expected, not only between species but within [1]. GenBank for example currently holds over 10000 genome assemblies for *E. coli*, one of the prokaryotic model organisms, displaying stunning diversity. The vast number of sequenced prokaryotes, across all different phyla, also makes it impractical to perform genome comparison based on sequence alignments to unravel the genomic complexity [2]. This makes novel *in silico* methods for genome annotations necessary.

The delineation of the open reading frame (ORF) is an essential element in gene annotation and is mostly performed *in silico* [3, 4]. Recently, ribosome profiling (also called ribo-seq) was introduced, measuring mRNA that is associated with ribosomes by sequencing ribosome-protected fragments (RPFs) [5, 6]. Ribo-seq experimentally enables the ORF delineation, and the technique has already been successfully adopted for prokaryotes [7, 8]. An important aspect of the ORF delineation is the determination of the Translation Initiation Site (TIS). Here also, specific prediction tools are in place to perform this task [9, 10, 11], but these TIS can also be detected by applying a specific antibiotic treatment (e.g. chloramphenicol or tetracycline) preceding the ribo-seq protocol enriching for initiating ribosomes [12]. Recently, prediction methods based on machine learning algorithms have been devised to either delineate the ORF [13] or predict the TIS [14] based on a combination of ribosome profiling and sequence features for prokaryotic genomes. A multitude of tools are also available for eukaryotic samples [15, 16, 17, 18, 19, 20].

Alternative proteoform usage can also be investigated by specific mass spectrometry protocols measuring N-terminal peptides [21, 22]. Although the technology is recognized, it suffers from drawbacks (e.g. peptide physical properties and modifications, mass spectrometry measurement range…), limiting the number of detectable N-termini. In order to attain a more comprehensive map of proteoform usage, proteogenomics studies have combined aforementioned high-throughput sequencing and mass spectrometry information, resulting in more precise ORF and TIS validation and thus genome annotation. [23, 24].

In this article we present DeepRibo, a novel neural network implementation applying ribosome profiling data for the precise annotation of TISs in prokaryotes. The use of artificial neural networks, which have proven to be highly effective on solving complex methods given the availability of sufficient data, is still confined to few applications in the field of bioinformatics. Examples are the use of convolutional neural networks for the prediction of DNA- or RNA-binding with a target protein [25] or precise variant calling on next-generation sequencing (BioRxiv: https://doi.org/10.1101/092890). DeepRibo is an artificial neural network that applies both convolutional neural network (CNN) and recurrent neural network (RNN) architectures in order to attain and combine information from the ribosome profiling signal and DNA sequence. To prevent biases introduced by sequence alignment techniques [26], only a short DNA sequence of 30 nucleotides covering the Shine-Dalgarno region is processed by the neural network.

DeepRibo is trained on a combination of available experiments for different bacteria and has been tested to work equally well on *de novo* ribo-seq data of bacterial genomes. We managed to successfully train a highly precise model that is able to process ribo-seq data without loss of resolution. We further validated our results with multiple species sequence similarity comparison [27], available mass spectrometry data and translation initiation site annotations [28].

DeepRibo is trained on data collected from ribosome profiling data. Ribo-seq data has the advantage that it does not map the untranslated regions of the transcribed mRNA. Ribosome profiling data upholds a high resolution and low background noise, making precise gene annotation possible. In prokaryotes, no splicing of the mRNA occurs, giving rise to more straightforward patterns of the signal along the coding regions as compared to eukaryotes. Conversely, bacterial genes are tightly packed and are frequently overlapping, which impedes a straightforward annotation. In order to detect genomic features, the model is designed to evaluate a set of possible ORFs containing ribo-seq signal, from which the top k probability scores are predicted to be expressed genes. The model is furthermore trained on a short DNA sequence covering the Shine-Dalgarno region, the ribosome binding site which has proven to be of major importance in predicting the presence of a TIS [9, 29]. Since DeepRibo has been designed to learn mainly from experimental data, the DNA sequence taken into account covers only 30 nucleotides, spanning from 20 nucleotides upstream to 10 nucleotides downstream of the considered (near-cognate) start triplet.

**Figure 1:**
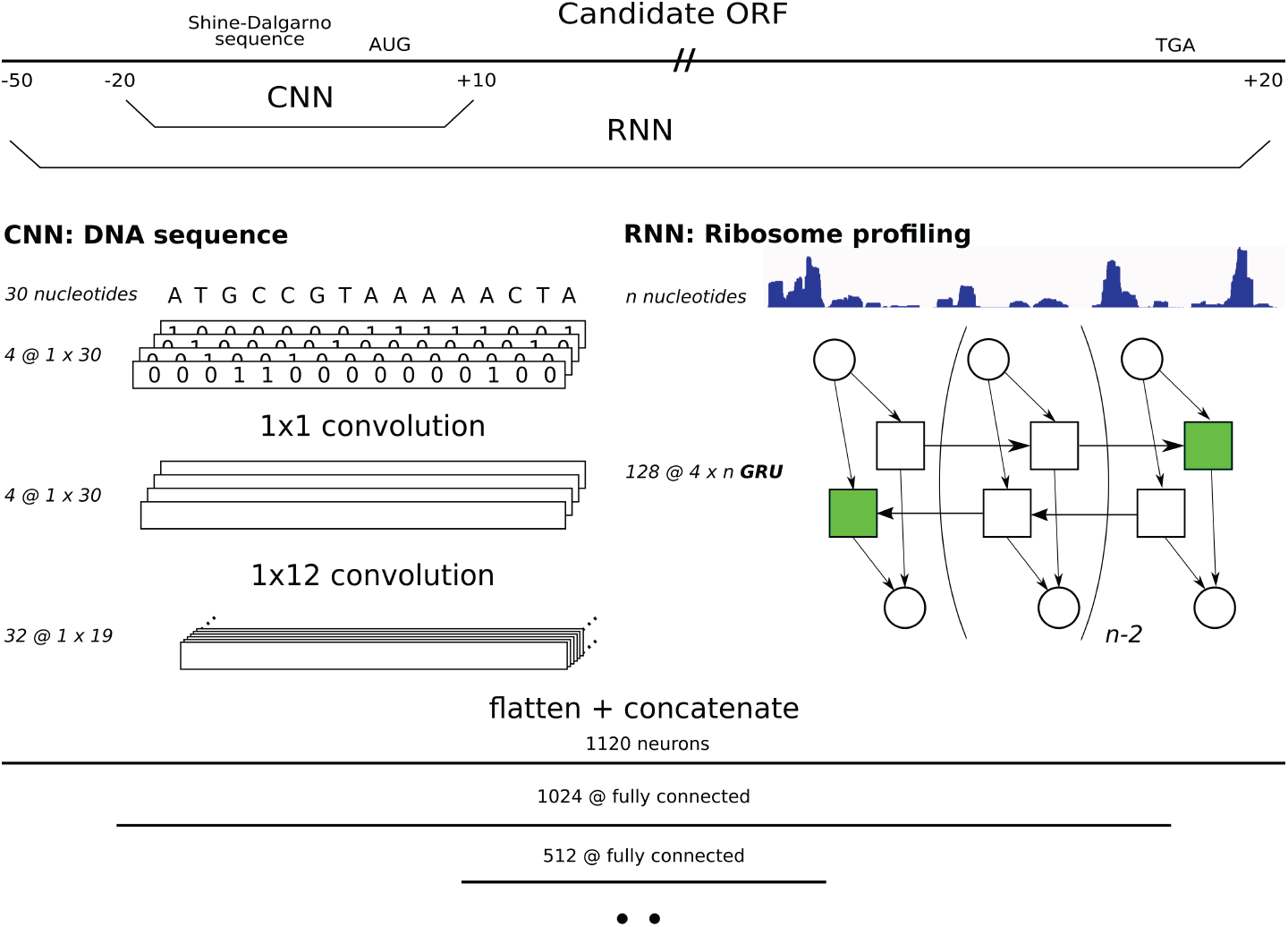
The architecture of the neural network DeepRibo. For each candidate ORF two types of data are processed and fed into their respective parts of the neural network. The convolutional layers train on a 30 nucleotide DNA sequence ranging from 20 nucleotides upstream to 10 nucleotides downstream of the TIS. The recurrent neural network covers the complete ORF from 50 nucleotides upstream of the start codon, including the SD region, and extending 20 nucleotides downstream of the stop codon. The DNA sequence is first translated in a binary image before being processed by four 1×1 and 32 1×12 convolutional kernels, respectively. The ribosome profiling data is processed by a double layered bidirectional GRU containing 128 weights. The outputs of both neural networks are flattened and concatenated and fed into three consecutive fully-connected layers of length 1024, 512 and 2.

### 1.1. Sample selection using the four parameter S-curve

The input ORF samples, labeled using the latest genome assemblies of the species, is the collection of all possible ORFs meeting a minimum signal strength. Since ribosome profiling changes according to the expression profile of the organism at the time of the experiment, no signal is present along many parts of the genome. Practically, it is not possible to make any predictions about these regions based on the ribosome profiling signal. Before selection of the positive and negative data, all candidate ORFs containing low ribosome profiling signal are therefore not considered when training/evaluating the model. The remaining data is afterwards labeled using the assembly files retrieved from NCBI. Other approaches including all possible ORFs have not been taken as bias was believed to be introduced. Indeed, leaving in positively labeled data with low to missing signal conditions the model to positively label candidate ORFs, based solely on the short DNA sequence. In contrast, negatively labeling a selection of the positive set, which does not meet the required minimum cutoff, biases the model towards signal strength across the different samples. The selection of data is based upon two properties of the samples, the coverage and signal read count. The coverage indicates the nucleotide fraction of the candidate ORF at which signal is present. The signal read count, expressed as Reads Per Kilobase Million (RPKM), expresses the amount of reads within the sample as compared to the dataset read count. Since the biggest partition of the considered dataset has zero to low coverage and RPKM, a more balanced distribution of read count, coverage, and label values is obtained from the filtered input samples. Moreover, the final dataset contains about one-fifth of the input ORF samples as compared to all candidate ORFs present in the data collection.

To determine the minimum cut-off values for coverage and RPKM, a method introduced by Ndah et. al. [13] has been applied. The method is based upon threshold dose-response estimation done by [30]. For this, a four parameter S-curve is fitted on the coverage in function of the RPKM. Only the positive samples are considered when fitting the S-curve. By predicting the lower bend of the fitted S-curve, minimum cut-off values of the signal coverage and RPKM for each dataset are obtained. Intuitively, this point is of importance as it separates the point at which positive samples can be distinguished from the background noise. This can also be defined as the point from which an increase in RPKM within the positively labeled candidate ORFs is correlated to the coverage of the ribo-seq signal in said dataset. Using this technique, it is possible to pool the data from several individual datasets, as the S-curve is fitted on each experiment individually.

To label the samples, the public genome annotations of the referred species are used. Indeed, the assumption is made that DeepRibo, trained on data labeled via sequence alignment, can offer precise predictions by learning from the ribo-seq signal instead of using the full DNA sequences as an input. Although it is expected that the annotated genomes contain errors because of the shortcomings of prevalent but more conservative DNA sequence alignment methods, this behaviour is not mimicked as the model does not learn the DNA sequences of the coding sequences.

### 1.2. Neural network architecture

DeepRibo is a neural network built in PyTorch [31]. It is specifically designed to process two types of data: strings (i.e. DNA sequences) and floats (i.e. ribo-seq signal). The model first processes each type of data in parallel before combining the features created from both inputs into a set of fully-connected layers. The DNA sequence is transformed into a binary image with 4 channels, a method proposed by Alipanahi [25]. This image is consecutively processed by two convolutional layers. The first layer transforms the sparse matrix into a dense matrix using four 1×1 convolutional kernels. Afterwards, 32 kernels of 1×12 convolutions process the data in the second and last convolutional layer. The ribosome profiling data is fed into a double-layered, bidirectional Gated Recurrent Unit (GRU). The gated recurrent unit was selected instead of the long short-term memory cell as it showed to train better models and was overall faster to train. Only the final hidden states of the memory cell are used for further processing, making the use of varied length inputs (i.e. candidate ORFs) possible. After each type of data is processed, the output nodes of both networks are concatenated and fed into a fully-connected layer. The final layers of the network consist of three fully connected layers that combine the features of both the Convolutional Neural Network (CNN) and Recurrent Neural Network (RNN) to obtain a final prediction. The rectified linear unit is used as the activation function after all but the last step. The binary cross entropy is used as the loss function during training. Figure 1 features the full architecture of the neural network.

### 1.3. Dataset construction

Several databases have been included for training, consisting of experiments performed on prokaryotes grown under standard conditions. The experiments cover both gram-negative (*Salmonella typhimurium* [13], *Escherichia coli* [32], *Caulobacter crescentus* [33]) and gram-positive bacteria (*Bacillus Subtilis* [34], *Streptomyces coelicolor* [35], *Staphylococcus aureus* [36]). The model is trained using the coverage profiles of the ribosome profiling data. The S-curve is fitted on each dataset to obtain the minimum required coverage and RPKM signal of the ribosome profiling signal of the samples within each dataset. Table 1 gives an overview of the used datasets, and the amount of samples each contributes to the the training/test data.

To make sure no bias is introduced during the creation of the input data, the first step selects all candidate ORFs of the genome for each of the included ribo-seq datasets. It has been shown that ATG, GTG and TTG are the three nucleotide combinations that almost exclusively make up all start codons in a wide variety of bacteria [37]. Therefore, all DNA sequences within the genomes starting with either ATG, GTG and TTG up until a stop codon (TAA, TGA or TAA) are considered candidate ORFs. Since a large amount of ORFs exist with lengths too short to be translated into a functioning protein, a pseudo-arbitrary cutoff of 30 nucleotides is chosen to be the minimum length of the samples.

The study is built up as follows: the training data is created from five out of the six available datasets, using the remaining dataset as the test set. Six models have been trained for this study, using each of the available datasets as a test set. Furthermore, to investigate the predictions of the model in more depth, the predictions of two models have been extensively evaluated. In set-up 1 (S1) we exclude the expression data from *S. aureus* from the training set. In set-up 2 (S2) *E. coli* is excluded from the training set. These are both the smallest and biggest possible test set, furthermore having the lowest and highest correlation between RPKM and coverage of the annotated genes, respectively. All experiments evaluate the performance of DeepRibo on *de novo* data (i.e. transfer learning), in accordance to the design goals discussed. The different neural networks selected were trained for about 11 epochs (Supplementary Figure S3,S4), based upon the minimum loss of the model’s predictions upon the test sets (Supplementary Figure 3).

**Table 1:** Different input organisms that make up the final dataset used to train and validate DeepRibo. To obtain a more balanced distribution of the labels and RPKM, each dataset has been filtered by applying a minimum threshold on coverage and RPKM. Cut-off values have been determined by estimating the lower bend point of the fitted S-curve.

| Dataset | Original data |  | S-curve selection |  |
| --- | --- | --- | --- | --- |
|  | Negative set | Positive set | Negative set | Positive set |
| *S. typhimurium* [13] | 428,269 | 4,714 | 113,228 | 3,474 |
| *E. coli* [32] | 435,646 | 4,249 | 115,011 | 2,999 |
| *C. crescentus* [33] | 270,520 | 3,870 | 50,935 | 2,203 |
| *B. subtilis* [34] | 413,609 | 4,241 | 88,171 | 2,779 |
| *S. coelicolor* [35] | 540,068 | 7,746 | 26,518 | 1,359 |
| *S. aureus* [36] | 308,529 | 2,767 | 20,742 | 852 |
| Total | 2,396,641 | 27,587 | 414,605 | 13,666 |

### 1.4. Evaluation and post-processing

To evaluate the model, the Area Under the Precision-Recall Curve (PR-AUC) performance measure is used. Due to the highly imbalanced dataset towards the negative samples, a large change in false positives leads to only a small change in the false positive rate. As the eventual use of the model is focused on the prediction of the top k genes, PR-AUC is known to be a more informative measure [38]. Indeed, measured Area Under Receiver operating characteristic Curve (ROC AUC) values can be high even in cases in which the absolute amount of false positives (heavily) outweighs the absolute amount of true positives.

An important post-processing step of the model is focused on restricting the final predictions to a single start (SS) site within a region of two stop codons. The sequencing depth, reflected by the translation rates of the RNA, varies strongly between different gene regions. The assumption is made that prediction biases due to changing probability score distribution might exist between regions. These are introduced by differences in RPKM for candidate ORFs of one gene region as compared to another. Therefore, when determining a threshold to obtain the positive predictions, it occurs that multiple start sites are predicted within one region while not obtaining TISs in another region. With the assumption that every ORF has only one start site, only the highest prediction probability between two stop codons is of importance.

**Figure 2:**
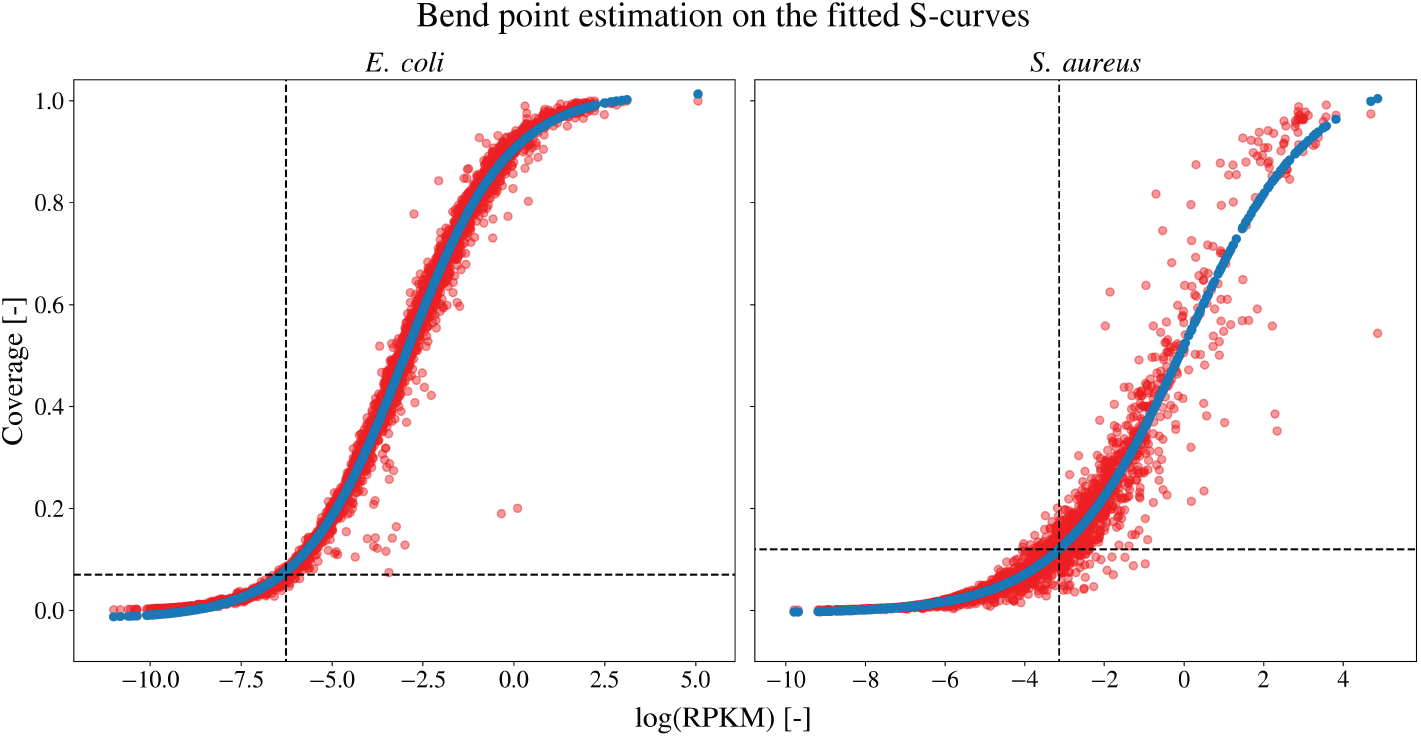
Bend point estimation on the fitted S-curves of the coverage in function of the log RPKM for both the *E. coli* (left) and *S. aureus* (right) dataset. The positive samples for each dataset (red) are plotted with the predicted (blue) ones for the fitted S-curve. For each dataset, the lower bend point of the fitted curve is estimated using the bent-cable function to obtain the minimum cut-off values.

### 1.5. Multiple sequence comparison based on local alignment

Given the performance measures for each of the models, a more in depth exploration of the results is made. Assuming the existence of mistakes in the annotation files, false positive predictions have been compared using the Basic Local Alignment Search Tool (BLAST) [27]. The false positive predictions of the model are compared to a database containing a collection of proteins which have been previously discussed in literature, forming a good criterion to evaluate the existence of the predicted ORF. A query of the false positive predictions on ‘the non-redundant protein sequences’ (containing non-redundant sequences from GenBank translations together with sequences from Refseq, PDB, SwissProt, PIR and PRF [39]) has been performed using protein-protein BLAST (pBLAST). A maximum cut-off value of 0.1 for the Expect (E) value is taken. The E value gives the expected amount of hits covering a similar alignment given the size of the database. For the sake of clarity, false postive predictions are considered as possible proteoforms or novel proteins, and are thus labeled as such. Specifically, proteoforms constitute false positive predictions with a varying start site compared to the positively labeled ORF. Novel proteins cover any false positive prediction for which no previous annotation was present.

**Figure 3:**
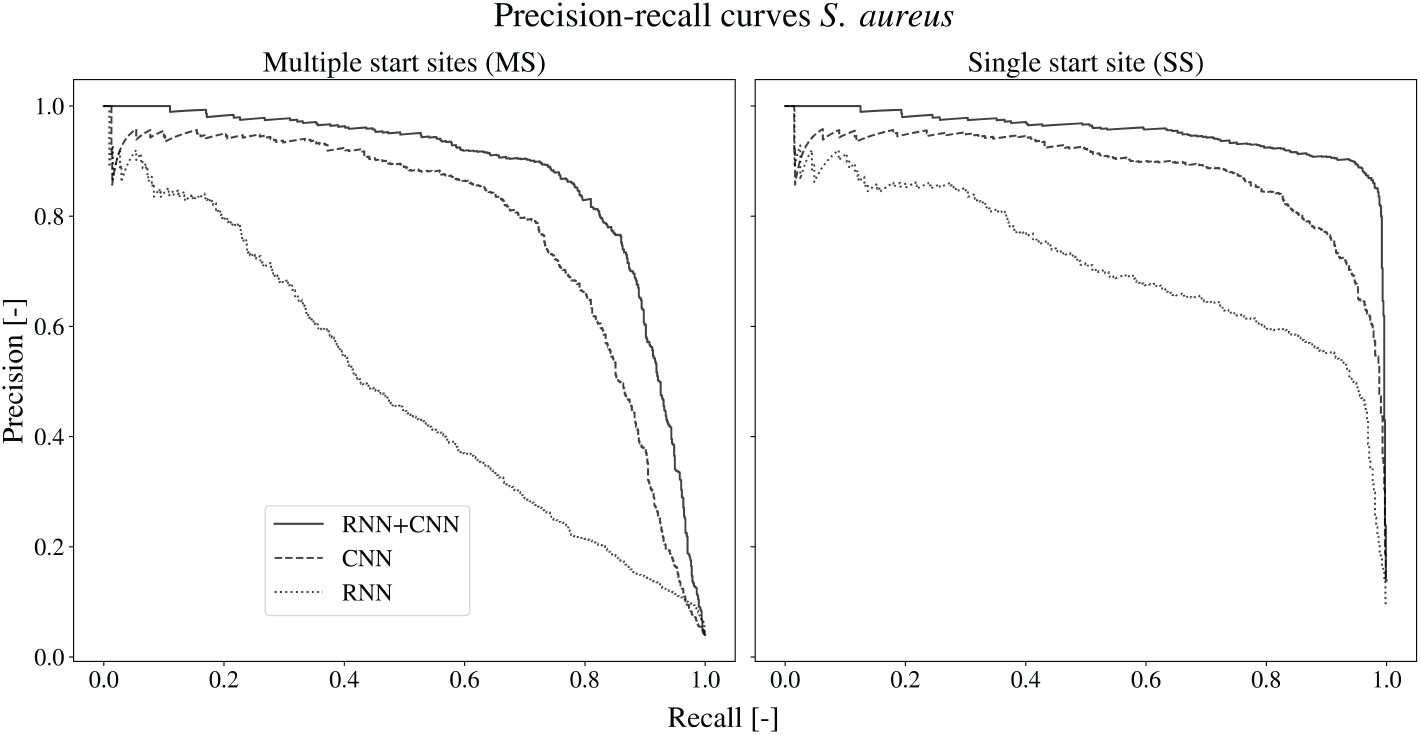
The precision-recall curves of the different networks on the *S. aureus* dataset. the precision-recall curves are given in case of the multiple start site and the single start site set-up. The full model (full line), combining the RNN and CNN outperforms both the single CNN (dashed) and RNN (dotted) architecture.

## 2. RESULTS

### 2.1. S-curve estimation for cut-off values filters high-quality from low-quality data

To normalize the total signal counts between multiple datasets, the expression rates of the different experiments are assumed to be equal. Since we are not working with repetitions of the same experiment, no normalization is performed before merging the datasets. However, differences in overall signal strengths between different experiments can be caused either by differences in expression profiles of the organisms, varying growth conditions, or technical variance introduced when performing the study. To filter ORFs with signal strengths indistinguishable from the background noise, minimum cut-off values are estimated for each dataset using the S-curve methodology [30] (Supplementary Figure 1). Interestingly, datasets containing a high amount of low expression values give rise to more stringent cut-off values (e.g. *S. aureus*). In case of a clear distinction between expressed and non-expressed genes, a relatively low cut-off value is obtained (e.g. *E. coli*). Therefore, depending on the quality of the data, the amount of samples selected from each dataset can vary greatly. The positive samples and the fitted S-curves for the *E. coli* and *S. aureus* dataset are plotted in Figure 2. In the case of an incorrectly annotated dataset, a decreased correlation between the coverage and RPKM of the positive samples is expected, with a shift of the data points towards the bottom-left corner. As these elements create a more gradual fit of the lower bend point of the S-curve on the data, these estimated cut-off values will be higher.

#### High performance values for predictions in the context of both single and multiple start codons

For the purpose of evaluating the performance, the test set is filtered to exclude any positively labeled data with low expression rates. As these genes are not being expressed, positive samples with non-existent or low ribo-seq data are filtered out (see Table 1). In parallel with the selection of the training set, minimum cut-off values have been determined using the fitted S-curve. Table 2 shows the performances of all the models on the independent dataset. Even though DeepRibo is trained using a maximum of one positively labeled ORF within two stop codons, this is not reflected into the predictions of the model. As genome assemblies are annotated using a maximum of one start codon for each stop codon, AUC and PR-AUC scores are consistently better including only the highest predicted start site for each stop codon. The performance of the model varies only slightly between the different experimental set-ups. A PR-AUC as high as 0.953 and 0.955 on the test set is obtained for S1 and S2, respectively. Although the existence of multiple start sites within prokaryotes has been confirmed [12], it can be expected that the predictions have shifted distributions between different regions due to a varying ribo-seq signal. However, even when considering multiple start sites, PR-AUC scores reach up to 0.880.

#### DeepRibo successfully combines DNA-sequence information and ribosome profiling data to optimize its performance

To confirm that the neural network is able to use the ribosome profiling data to make better predictions, two custom models have been trained on either the Shine-Dalgarno sequence (based on CNN) or the ribosome profiling data (based on RNN). The architectures of the models are kept similar, except for the loss of the recurrent or convolutional section, in case of the model trained on the DNA sequence and ribo-seq data, respectively. Table 2 lists the performances of both models using set-up 1. Both approaches prove to be effective at training from their specific data, with AUC values of 0.968 for RNN and 0.984 for CNN. The CNN has overall better performances than the RNN, especially when comparing its PR-AUC measure of 0.884 compared to 0.717 of the RNN. However, due to the exclusion of candidate ORFs with low signal, the data fed into the CNN is biased based on information it could otherwise not process. An important conclusion that can be made is that the combination of both neural networks works better compared to the individual parts. With an improvement of the PR-AUC of about seven percent compared to the CNN and 23 percent compared to the RNN, the model shows to be able to combine both types of information in a meaningful way.

#### Comparing the predicted and annotated ORFs reveals they predominantly disagree on TISs with less common start sites

Comparing the results of our model with the annotated dataset, an evaluation is made on the predicted ORFs. To obtain a set amount of positive predictions, a threshold is set on the output probabilities. After filtering the dataset from samples with insufficient signal or coverage, a final set of positive labels is obtained for each dataset. The resulting total amount of positive labels can be used as a conservative estimation on the expected amount of genes expressed in the relevant organism. Accordingly, a threshold is set to obtain that amount of positive predictions. However, depending on the aim of the study, custom methods can be applied to set the threshold for the final results. The precision of the top 852 positive predictions for the model in S1 (see Table 1) is 0.862, having 118 false positives. With 365 false positives the precision of the top 2,999 predicted TISs for the model in S2 (see Table 1) is 0.878.

**Table 2:**
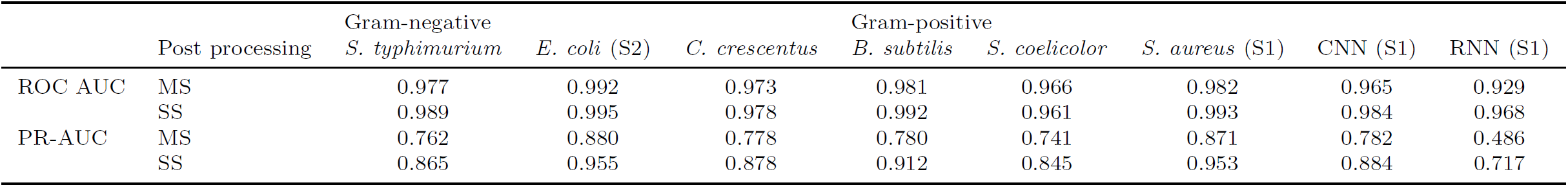
The ROC AUC and PR-AUC performance values for the different experimental set-ups in which each of the available datasets is excluded from training. The performance metrics for are given in case multiple start sites are considered possible (MS) and in case each stop codon can only have a single predicted start site (SS). The last two columns list the performances of the model tested on the smallest dataset *S. aureus* (SI) and containing only the CNN and RNN architectures.

**Table 3:** Start codon frequencies amongst the positive set. Start codon prevalence is summarized for three groups, the True Positive (TP), False Negative (FN) and False Positve (FP) results.

| Start codon | *S1 (S. aureus)* |  |  | *S2 (E. coli)* |  |  |
| --- | --- | --- | --- | --- | --- | --- |
|  | TP | FN | FP | TP | FN | FP |
| ATG | 0.924 | 0.796 | 0.592 | 0.911 | 0.805 | 0.814 |
| GTG | 0.047 | 0.117 | 0.287 | 0.070 | 0.157 | 0.123 |
| TTG | 0.029 | 0.087 | 0.121 | 0.019 | 0.038 | 0.063 |

**Figure 4:**
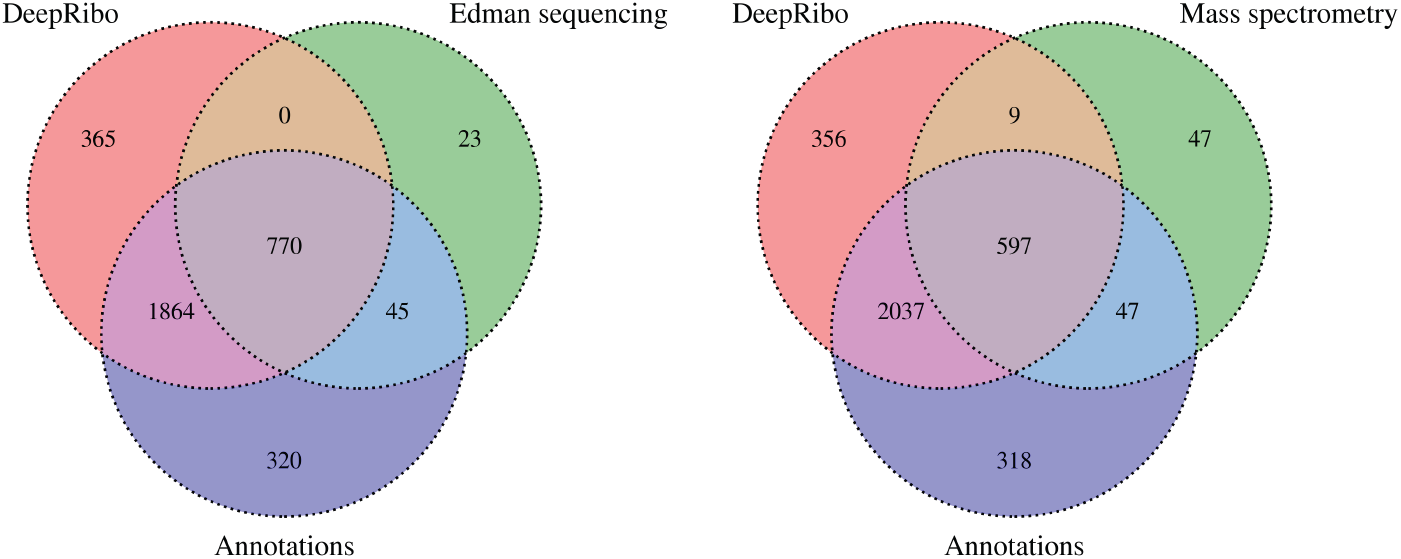
Venn diagram displaying the distributions of the proteins verified by Edman sequencing (left) and mass spectrometry (right) within the annotated dataset and model predictions. Distributions only include expressed ORFs, determined using the S-curve methodology.

Table 3 gives a representation of the nucleotide use along the TISs of the true positives, false negatives and false positives. A certain shift can be observed between the start codon use of the true positives, false negatives and false positives. The results seem to indicate that the model has higher difficulty correctly annotating TISs for less common start codons.

### 2.2. Edman degradation assisted validation of predictions

Through sequencing of the N-terminal residues of the matured proteome using Edman degradation, the creation of certain proteins within a cell can be verified. A collection of 922 proteins within E.coli K-12, featuring all the verified proteins discussed in literature, is featured by Ecogene [28]. Of the 922 proteins, a total amount of 838 ORF are expressed within the *E. coli* dataset, determined using the S-curve methodology. The positive predictions are composed of the top 2999 predictions, using the single start site setting, in accordance to previous methods. 770 (91.9%) of the genes have been predicted correctly by the model. 23 (2.7%) verified proteins have TISs differing from the annotation, resulting in 815 proteins for which the annotation and verified protein set agree. None of the predicted TISs in agreement with the verified proteins were in disagreement with the labeled dataset. 45 out of 815 (5.5%) TISs present in the annotations and Ecogene dataset are not picked up by the model. However, 15 out of the 45 false negatives are the predicted TISs within the single start site setting, and are actually all present in the top 4000 predictions. This means only 30 out of 815 (3.7%) of the false negatives have predicted TISs up- or downstream of the annotated gene. Due to the inclusion of novel ORF within the model’s predictions, some of the annotated regions are bound to be excluded from the positive predictions when setting the threshold to obtain a number of positive predictions equal to the amount of positive samples.

### 2.3. N-terminal proteomics based validation of predictions

Next to the Edman sequencing (Ecogene dataset), mass spectrometry based proteomics can also serve to validate our predictions. N-terminal proteomics, more specifically, is a technology that enables us to detect N-terminal peptides compliant with the rules of initiator methionine processing. 781 such N-termini were previously determined for *E. coli* [13]. 700 N-terminal peptide sequences that are aligned with coding sequences are expressed and are therefore present in the test set. 644 out of 700 samples (92%) are in accordance with the annotation. 47 (7.2%) of the 644 samples are not predicted by the model, of which 32 have differing TISs and 15 fell out of the top 2999 predictions. Interestingly, of the 56 peptide sequences which indicate a TIS in disagreement with the annotation, 9 have been predicted by DeepRibo. Although the presence of a TIS at a site differing from the annotation can be suggested as indicated by the ribosome profiling data, this is tangible proof that the annotation is not waterproof, negatively influencing the performance measure of the model. Figure 4 gives an overview of the overlap between the two validation datasets with the annotations and predictions.

### 2.4. A high percentage of the false positives have highly identical matches in the non-redundant protein database

Multiple proteoforms exist for a large amount of the annotated proteins. Yet, only one variety of each protein has been annotated in the genome assembly. Biological variation, growth condition or growth phase are some of the factors influencing protein expression rates. Accordingly, variety in protein expression between different experiments creates variation from the annotated genome. pBLAST searches have been performed to investigate whether false positive predictions could be caused by expressed proteoforms not present in the annotation. A summary is created by simply taking the best aligned protein for each of the false positive predictions. pBLAST searches have been performed on the complete set of false positives for S1 and S2. The validation of the predictions using the mass spectrometry and Edman sequencing dataset resulted in 32 and 30 ORFs with differing TISs as proposed by the model. two sets of samples containing the alternative predictions of the model have furthermore been included for sequence similarity comparison. Table 4 gives an overview of the results. As expected, all proteoforms have been successfully aligned, given they are partly identical to the annotated gene. As much as 76 out of 86 (88.4%) and 179 out of 209 (85.6%) proteins of the proteoforms have been aligned with an existing protein sharing both the TIS and stop site for set-up 1 and 2, respectively. Of all novel proteins, a lower percentage could be aligned on both the start and stop site, summing up to a total of 16 out of 32 (50%) and 53 out of 156 (34.0%) protein sequences for S1 and S2. Interestingly, a considerable percentage of the novel proteins are described as ‘hypothetical’. The model predictions going against the results listed in the MS and Ecogene dataset mostly indicate perfect alignment with proteins present in the non-redundant database, with 27 out of 32 (84.3%) and 30 out of 32 (93.7%) matches, respectively. Figure 5 gives the spread of the E values for each of the aligned proteins. A complete list of the false positive and false negative predictions for *E. coli* and *S. aureus*, including the two validation datasets and the BLAST results is provided in Supplementary File 1.

**Figure 5:**
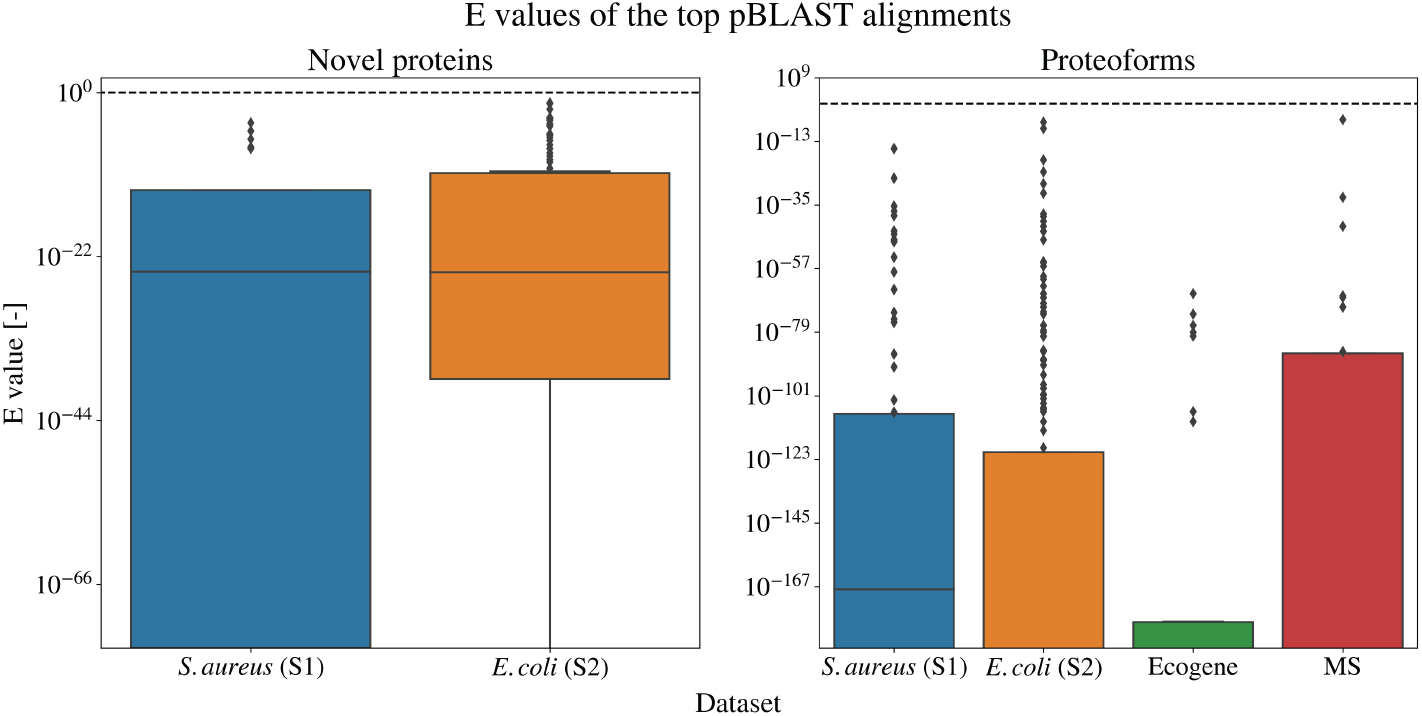
E value distributions for the pBLAST results on newly predicted proteins (left) and proteoforms (right) for the different datasets. The E values are given for the best hit (if existent) for each of the false positives. The dashed line indicates the E value of 1.

**Figure 6:**
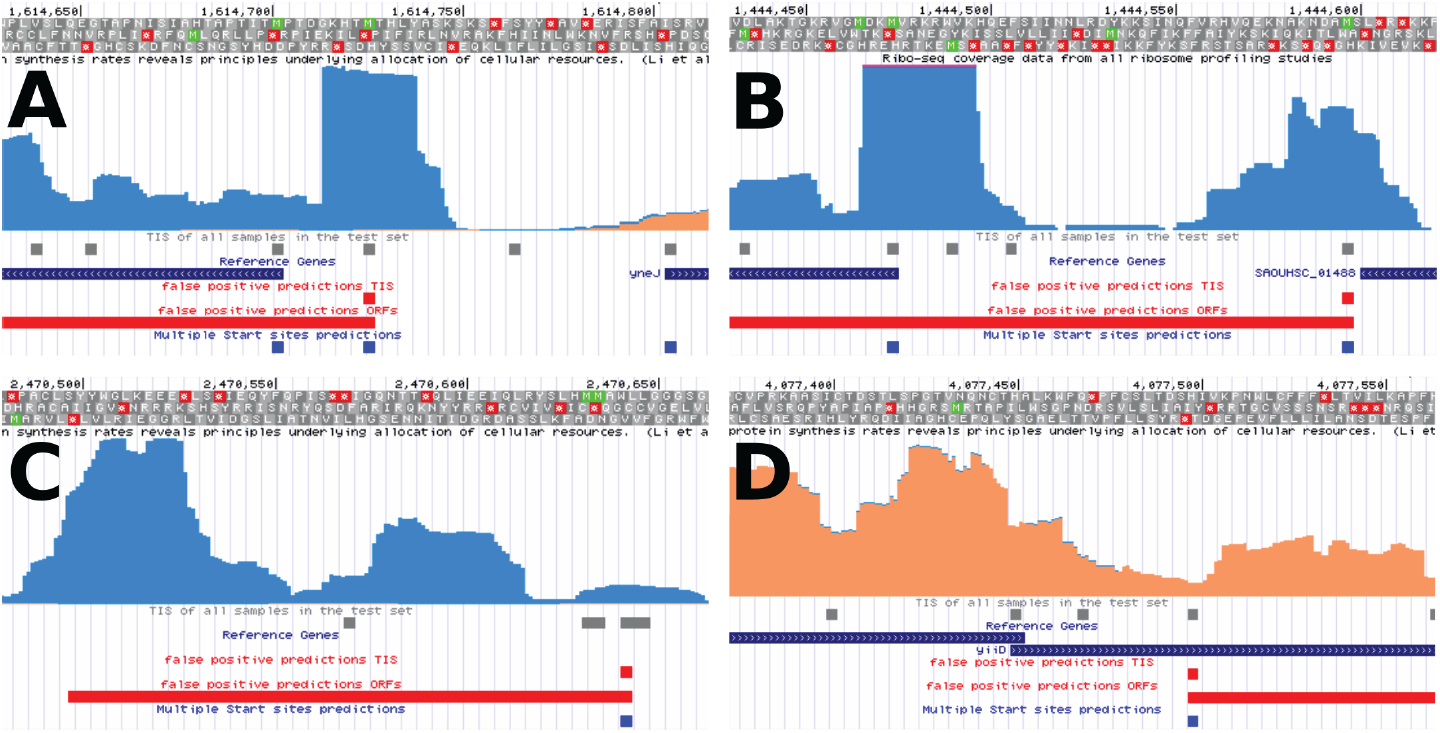
DeepRibo example predictions displayed alongside the ribo-seq input signal and the annotations. The data is formatted using the GWIPS-viz browser [40] and is hosted publicly (see Supplementary Data). On every track is displayed (from top to bottom): nucleotide position, amino-acid translations, ribo-seq signal (sense: orange, antisense: blue), TIS of all samples present in the test set, genome annotation, TIS of the false positive predictions, ORF of the false positive predictions, TIS of the predictions in a multiple start site setting. (A) The highest ranking proteoform prediction (rank: 127) for *E. coli*; (B) The highest ranking proteoform prediction (rank: 94) for *S. aureus*; (C) The highest ranking novel protein for *E. coli* with no pBLAST alignments (rank: 1470); (D) An example of a false positive prediction in a region with overlapping genes (rank: 941).

**Table 4:** Results of the BLAST search on the false positive set of the two set-ups (S1,S2), and specifically on the false positives directly going against results gained from Mass Spectrometry (MS) and Edman sequencing (Ecogene) dataset. These predictions can be divided into proteoforms, which have a TIS which is either up- or downstream of the annotated one, or novel proteins, constituting ORFs with a non-annotated stop site. A BLAST search of these proteins was performed on the non-redundant protein database. A maximum cut-off value of 0.1 for the E score is taken. The total amount of false positives are given for each type. Taking only the best aligned protein (i.e. highest E score) for each of the false positive results, the total amount of matches that were aligned by start site or both start and stop site are given. Finally, the total amount of proteins described as ‘hypothetical’ are also displayed.

| Set-up | type | # | aligned<br>total | TIS | TIS + stop | description<br>hypothetical |
| --- | --- | --- | --- | --- | --- | --- |
| **S1** | Proteoform | 86 | 86 | 85 | 76 | 11 |
|  | Novel protein | 32 | 21 | 18 | 16 | 6 |
| **S2** | Proteoforms | 209 | 209 | 192 | 179 | 32 |
|  | Novel protein | 156 | 91 | 58 | 53 | 47 |
| **MS** | Proteoforms | 32 | 32 | 27 | 27 | 0 |
| **Ecogene** | Proteoforms | 30 | 30 | 27 | 27 | 3 |

## 3. DISCUSSION

The success of deep learning methods on popular topics involving big data is slowly finding its way to the field of bioinformatics involving multi-omics. Although big data created by high-throughput methods has been available since the arrival of second generation sequencing, it has so far mainly been explored using statistical methods, excluding machine learning. Deep learning has proven to be considerably successful, allowing the use of a black box approach when interpretability is not important or feasible. In this study, we present a deep neural network for the precise annotation of expressed proteins on the genome using ribosome profiling data. This tool combines data from an *in vivo* experiment to increase the accuracy of *in silico* based methods. DeepRibo learns from information contained in both DNA sequences and ribo-seq, using a novel architecture which combines both convolutional layers and recurrent memory cells. Results obtained from machine learning models, which are trained and evaluated on the same dataset, can be overestimates of their performance on new data due to overfitting. The use of a single model trained on a variety of existing datasets and evaluated on independent test sets makes due with this problem. Moreover, building the model on a combination of datasets trains it to differentiate between useful features present over all the datasets and dataset-dependent variations, possibly making the need for normalization steps redundant. DeepRibo is the first tool for the precise delineation of ORFs in prokaryotes trained and validated on multiple datasets. It furthermore outperforms REPARATION [13], reporting PR-AUC values of 0.74, 0.80 and 0.89 after 10-fold cross validation when trained and tested on the *S. typhimurium, E. coli* and *B. subtilis* dataset, respectively.

The performance of DeepRibo is consistent on all six test sets, with a difference of 0.11 in PR-AUC score between the best and worst performing model. No difference is observed on the performance between gram-positive and gramnegative bacteria. Discrepancies between the predictions and the annotation are present and can be attributed to several factors discussed.

When evaluating the results of DeepRibo, a certain cut-off has to be determined to determine the fraction of positive predictions. To evaluate the model, the amount of positive ORFs has been set equal to the ORFs present in the annotations. However, due to novel predictions being made, a fraction of the annotated samples are bound to have a rank lower than the top k predictions (especially in a multiple start site setting). This is furthermore reflected by the fraction of proteins in the validation sets not picked up by the top k predictions of the model. No cut-off is optimal for every instance and has to be determined in line with the application, which postulates the desired precision/recall.

Since the majority of the candidate ORFs share their stop sites with other samples, selecting only the ORF with the highest predicted probabilities within each group gives consistently better performances. Even though an increase in performance is observable when comparing the single start site with the multiple start site setting, the performance of the latter is still noteworthy. Specifically, 105595 out of the 113228 (89.5%) candidate ORFs share stop sites with other samples in the *E. coli* dataset. Some regions have as much as one hundred possible TISs. Although the model has no way of processing this information, making a prediction on every sample individually, it achieves remarkable PR-AUC scores on the test sets, ranging from 0.762 to 0.880 on all models. Part of this error is expected to be caused by differences in RPKM values existent between different genome regions. Yet, the models’ performances indicate this effect to be minimal. Moreover, recent studies have discovered genes with multiple translation initiation sites [41, 14, 12]. As this feature is not supported by current annotations, correct evaluation of the model in a multiple stop site setting is not possible.

In case of the *E. coli* model, many of the novel predictions are situated within a pseudogene. Typically, no candidate ORFs overlapping the complete pseudogene regions were present in the training/testing samples, as these annotated features cover regions with multiple stop codons. Therefore, no positively labeled samples are present. However, ribo-seq signal is often measured at these sites, creating a hot-spot for ‘novel’ (false positive) predictions.

The identification of a high amount of novel small open reading frames (sORFs) by the model presents another contrast with the annotation. The novel ORF predictions given by the models have a median length of 165 and 63 for *E. coli* and *S. aureus*. These are well below the median length of the annotated genes within each species (807 and 723). The size of the ORFs influences the power of the statistical methods used for the identification of the sORFs by *in silico* methods [42]. A higher amount of sORFs are expected to be present on the genome than given by the annotation, several of which might be picked up by DeepRibo. An example of a novel prediction for *E. coli* is given in Figure 6C.

Many prokaryotic systems have a closely knit operon structures [43], creating a ribo-seq signal which can be overlapping over different regions of interest. Evaluation of the predictions has shown DeepRibo to have multiple false positives around these regions (Figure 6D). By inclusion of a padded region around the ribosome profiling signal processed by the RNN, DeepRibo showed increased performance. This is likely due to the possibility of the model to differentiate between samples laying within expressed genes and those for which lower or no ribosome profiling signal exists at their borders. However, genes closely located to each other can share similar patterns at their borders to candidate ORFs located within expressed regions, tricking the model into making wrong predictions.

Corroborated by the results obtained from the pBLAST searches, it is likely that a fraction of the false positives observed when evaluating the predictions of the single start site setting are due to an annotation which does not fully map the translational complexity of the organisms. Moreover, detailed evaluation of the predictions with the ribo-seq signal shows that many false positive results can be explained by the signal (Figure 6A/B/C).

As DeepRibo is a neural network that has been trained on only six datasets, it is safe to assume that performances will increase as more data for training is included. A more complete annotation of the assemblies on which the model is trained is expected to further improve DeepRibo.

## 4. SUPPLEMENTARY DATA

A list containing all samples, its metadata and the predictions given by the model for each of the test sets/models is featured in Supplementary File 1. Supplementary File 2 contains specific sets of samples as discussed in the article. Supplementary File 3 contains supplementary figures and information. For each model, predictions have been visualized and displayed alongside the input signal using the GWIPS-viz browser (Figure 6), accessible at (http://www.kermit.ugent.be/files/gwips_hub/index.html). A python module for DeepRibo is hosted at the BIOBIX GitHub repository (https://github.com/Biobix/DeepRibo).

## 5. ACKNOWLEDGEMENTS

The authors acknowledge the support of Ghent University.

